# Proteomic Signatures of Hepatitis B Virus Mutations Reveal Genotype-Specific Host Responses and Biomarker Candidates

**DOI:** 10.1101/2025.09.04.674240

**Authors:** Ahrum Son, Eun Ju Cho, Jaeho Ji, Ziyun Kim, Su Jong Yu, Bum-Joon Kim, Hyunsoo Kim

## Abstract

Hepatitis B virus (HBV) remains a global health challenge, with viral genetic heterogeneity and mutation-driven resistance complicating treatment outcomes. While previous genomic and transcriptomic studies have characterized HBV mutations, the proteomic consequences of these variants remain underexplored. In this study, we applied liquid chromatography– mass spectrometry (LC–MS)–based proteomics and systems biology approaches to serum samples from 60 HBV-infected patients, stratified by mutation-defined genotype signatures. Four genotype groups were generated, including those harboring mutation related to liver disease progression (rt269/s184) and basal core promoter/precore mutations (A1762T/G1896A). Comparative analyses of 406 high-abundance plasma proteins revealed distinct proteomic signatures, particularly in the ##AG group lacking mutation related to liver disease progression motifs. This group exhibited elevated CRISPLD2 and HSPD1 expression, implicating a dual axis of anti-inflammatory buffering and chaperone-mediated viral processes. Network and co-expression analyses identified modules enriched in focal adhesion, extracellular matrix remodeling, angiogenesis, and PI3K/Akt signaling—pathways tightly linked to hepatocarcinogenesis. Protein–protein interaction enrichment further highlighted disruption of chaperone networks, cytoskeletal regulation, and unfolded protein response. These findings provide molecular evidence for genotype-specific host–virus interactions, nominate CRISPLD2 and HSPD1 as biomarker candidates, and suggest therapeutic strategies targeting PI3K/Akt and microenvironmental pathways. Our results underscore the value of proteomics in refining genotype-informed risk stratification and personalized management in chronic HBV.

## Introduction

Hepatitis B virus (HBV) contributing to the development of cirrhosis and hepatocellular carcinoma (HCC), remains a major health burden with 250 million people chronically infected worldwide (1). HBV infection follows a complex lifecycle involving host cell entry, hijacking of cellular machinery for viral replication, establishing persistent infection through immune evasion mechanisms and integration into the host genome that lead to chronic inflammation and progressive liver damage. The heterogeneity of HBV was contributed by viral load and host immune status, but also by specific viral mutations that that modulate replication capacity, transcriptional activity, and immune evasion(2). The transition from acute to chronic infection, and subsequently to cirrhosis and HCC, involves intricate virus-host interactions that vary significantly among patients, with current treatment options including nucleos(t)ide analogues and interferon therapy achieving functional cure in only a small percentage of patients, largely due to viral heterogeneity and mutation-driven resistance.

Naturally occurring mutations in the reverse transcriptase (rt269LI), in the surface gene (s184V/A), and in the basal core promoter/precore region (A1762T, G1896A) have been associated with altered viral phenotype, disease progression (3-11). Mutations in the reverse transcriptase are known to enhance polymerase activity and viral replication efficiency, often emerging under selective pressure from antiviral therapy (12, 13). The surface mutation has been linked changes in hepatitis B surface antigen (HBsAg) folding and induction of endoplasmic reticulum (ER) stress pathways, leading to increased virion production (14, 15). Mutations in the basal core promoter and precore region are related to the transcriptional level and reshaping viral gene expression programs, particularly affecting hepatitis B e antigen (HBeAg) production and core protein synthesis (16-18).The emergence of these drug-resistant mutations poses significant clinical challenges and underscores the need for personalized therapeutic approaches based on viral genotype and mutation profiles.

While underlying detailed mechanism following virus infection have been studied through transcriptomic and genomic approaches, how such mutation group shape the global host proteomic changes has not been systematically investigated. Previous studies have primarily focused on individual mutations or single pathway analyses, lacking the comprehensive systems-level approach necessary to understand the global impact of viral genetic diversity on host biology, with most proteomic studies in HBV examining advanced disease stages and providing limited investigation of how early mutation-driven changes influence disease trajectory (19, 20). Proteomics offers distinct advantages over genomic studies by capturing post-translational modifications, protein interactions, and functional pathway activities that directly reflect cellular phenotypes (21-23). Circulating proteins provide a valuable window into the molecular interplay between HBV activity and host responses, and global proteomic profiling can reveal pathways that reflect viral mutation-specific host consequences. Understanding mutation-specific host responses has critical clinical implications for risk stratification, treatment selection, and monitoring of disease progression, as current clinical markers including HBV DNA levels and liver enzymes provide limited insight into the complex molecular mechanisms driving disease heterogeneity among patients with different viral mutation profiles.

Here, we hypothesize that mutation related to liver disease progression and transcriptional reprogramming mutations induce distinct host proteomic signatures that reflect different biological functions. Using liquid chromatography-mass spectrometry (LC-MS) based proteomics and systems biology approaches, we analyzed blood samples from HBV patients with various mutation patterns to identify co-regulated protein modules associated with different genotypes of HBV virus and characterize the biological pathways preferentially modulated by each mutation category. Our findings provide important insights for personalized medicine approaches in HBV management and advance our understanding of virus-host interactions at the systems level.

## Materials and Methods

### Sample Preparation

From July 2013 to December 2022, serum samples were obtained from 60 patients with chronic HBV infection at Seoul National University Hospital. The cohort consisted of 24 patients diagnosed with chronic hepatitis B and 36 with liver cirrhosis, of whom 33 were also diagnosed with HCC. Among all samples, 52 were HBeAg-negative and 8 were HBeAg-positive. At the time of enrollment, 35 patients received antiviral therapy with medications such as besifovir (BSV), lamivudine (LAM), adefovir (ADV), entecavir (ETV), or tenofovir (TDF, TAF).

The study was conducted in accordance with the guidelines of the Declaration of Helsinki and approved by the Institutional Review Board (IRB) of Seoul National University Hospital (IRB No. 2303-136-1414). Samples obtained from the Korea Biobank Network were collected with informed consent under IRB-approved protocols.

### DNA extraction, PCR amplification, and Sequences

HBV DNA was extracted from 200 μl of serum using the QIAamp DNA Blood Mini Kit (QIAGEN Inc., Hilden, Germany), dissolved in Tris-EDTA buffer, and stored at -20 °C for further analysis. To investigate mutation patterns and frequencies in the surface (S) region and PreC/C region, primers specific to the HBV genome were used. Since the S region completely overlaps with the reverse transcriptase (RT region), primers targeting the RT region and a nested PCR protocol were employed to sequence the entire S genome. The nested PCR and primer design followed previously established protocols (10). A nested PCR protocol was specifically applied to detect mutations at residue 184 within the S region. In the first round of PCR, the sense primer HB1F (5’-AAG CTC TGC TAG ATC CCA GAG T-3’) and the antisense primer HB3R (5’-AGT TGG CGA GAA AGT GAA AGC CTG-3’) were used. For the second round, the sense primer HB2F (5’-TGC TGC TAT GCC TCA TCT TC-3’) and the same antisense primer HB3R were utilized. This protocol was repeated for all samples to ensure consistent mutation detection. For the PreC/C region, direct PCR was performed using the sense primer HB7F (5’-GAG ACC ACC GTG AAC GCC CA-3’) and the antisense primer HB7R (5’-CCT GAG TGC TGT ATG GTG AGG-3’). Each PCR reaction was carried out in a 20 μl mixture containing 1.5 mmol/L MgCl2, 250 μmol/L dNTPs, and 1.0 U of ProFi Taq DNA polymerase (Bioneer, South Korea). The cycling conditions were as follows: initial denaturation at 95 °C for 10 minutes, followed by 30 cycles of 94 °C (15 seconds), 55 °C (15 seconds), and 68 °C (1 minute), with a final extension at 72 °C for 5 minutes. Across all samples, 68.8% yielded positive results. Publicly available HBV genome sequences of genotype C from a previous study (7) were included in the analysis. Among 892 sequences classified as genotype C2, 666 belonged to the s184V group, 226 to the s184A group, and 1 sequence was unclassified. Of these, 272 sequences were categorized as genotype C2(3), with 56 sequences in the s184V group and 216 sequences in the s184A group.

### Protein Extraction and Digestion

Serum samples were prepared for LC-MS/MS-based quantitative proteomic analysis by enzymatic digestion. Each tube, containing 100 μg of protein, was first treated to reduce disulfide bonds using 2% sodium deoxycholate (SDC) and 10 mM tris-(2-carboxyethyl)-phosphine hydrochloride (TCEP) at 60□ for 60 minutes. To block free cysteines, 20 mM iodoacetamide (IAA) was added and the samples were incubated in the d ark at 25□ for 30 minutes. The protein solution was then diluted to a final concentra tion of 20 mM ammonium bicarbonate (ABC) with distilled water. Trypsin (Promega, Sequencing Grade Modified, 20 μg) was added at a 1:50 enzyme-to-protein ratio and digestion proceeded overnight at 37□. The reaction was quenched with 1% formic a cid. Finally, 5 μL of a 6 × 5 LC-MS/MS Peptide Reference mix (Promega) was added to the digested samples, which were stored at -20□ until ready for analysis.

### Multiple Reaction Monitoring -Mass Spectrometry (MRM-MS) and MS data Analysis

The MRM-MS analysis was performed using an Agilent 1290 Infinity □ Bio 2D-LC system, coupled with an Agilent 6495C Triple Quadrupole mass spectrometer equipped with a Jet-stream electrospray ionization source. Peptide separation was achieved on a ZORBAX Eclipse Plus C18 column (Agilent, ID: 959759-902, 150 mm in length, 95 Å pore size, 1.8 μm particle size) maintained at 40□. The mobile phases used were 0.1% formic acid in water (Mobile Phase A) and 0.1% formic acid in acetonitrile (Mobile Phase B). Separation of the peptides from digested proteins was carried out over a 30-minute gradient, beginning at 3% Mobile Phase B for the first 4 minutes, increasing to 25% by 22 minutes, 40% by 24 minutes, and reaching 60% at 25 minutes. The system was then returned to 3% Mobile Phase B for re-equilibration at 28 minutes. The flow rate was maintained at 0.4 mL/min. Sample injections were made at a concentration of 1 μg/μL.

### MS data Analysis and Statistical Analysis

MRM-MS data were processed and analyzed using Skyline software (MacCoss Lab). For each target transition, the peak areas were normalized based on the chromatographic peak area from the 6 × 5 LC-MS/MS Peptide Reference Mix (Promega), matching the closest retention time and intensity. Statistical analysis was conducted in Python using the scipy library, applying a two-tailed unpaired Student’s t-test for comparison s between two independent groups. Fold-changes (FC) between groups were calculated in Excel (Microsoft Office 16) by dividing the intensity of one group by the intensity of the other.

### GO Analysis

Gene Ontology (GO) analysis was performed on the identified proteins using the DAVID platform. Statistical significance was assessed using the Benjamini and Hochberg False Discovery Rate (FDR) method. The analysis covered the categories of “biological process,” “cellular component,” and “molecular function”. GO analysis results were visualized using R software.

## Results

### Overall Study Design

In this study, blood samples (n=60) from patients with chronic HBV infection were used (Table 1). We focused on the 4 viral genes (rt269, s184, 1762, and 1869), and their specific genotypes for each gene; rt269L, s184V, 1762A, and 1896G. We stratified patients according to predefined genotype signatures in two functional blocks of the HBV genome. The first block genes comprised of the replication throughput enhancing (rt269, s184), in which the complete LV motif was considered as the signature. The second block of genes included the basal core promoter/precore region (A1762 an G1896), where the AG combination was defined as the signature. Variants were classified as “present” only when the entire block-specific signature was detected, whereas undetermined positions (“#”) were treated as missing. Based on these criteria, four mutually exclusive groups were generated: Group1 (LVAG), with both signatures present; Group2 (LV##), with replication through enhancing signature present only; Group3 (##AG), with basal core/precore region signature present only; and Group4 (####), with neither signature detected. Group 4 does not necessarily represent a wild-type group but rather a reference group with samples with undetermined genotypes at all signature positions. This grouping allowed evaluating the independent and combined effects of the replication throughout enhancing signatures and basal core/precore region signatures on host protein expression.

**Table 1.**
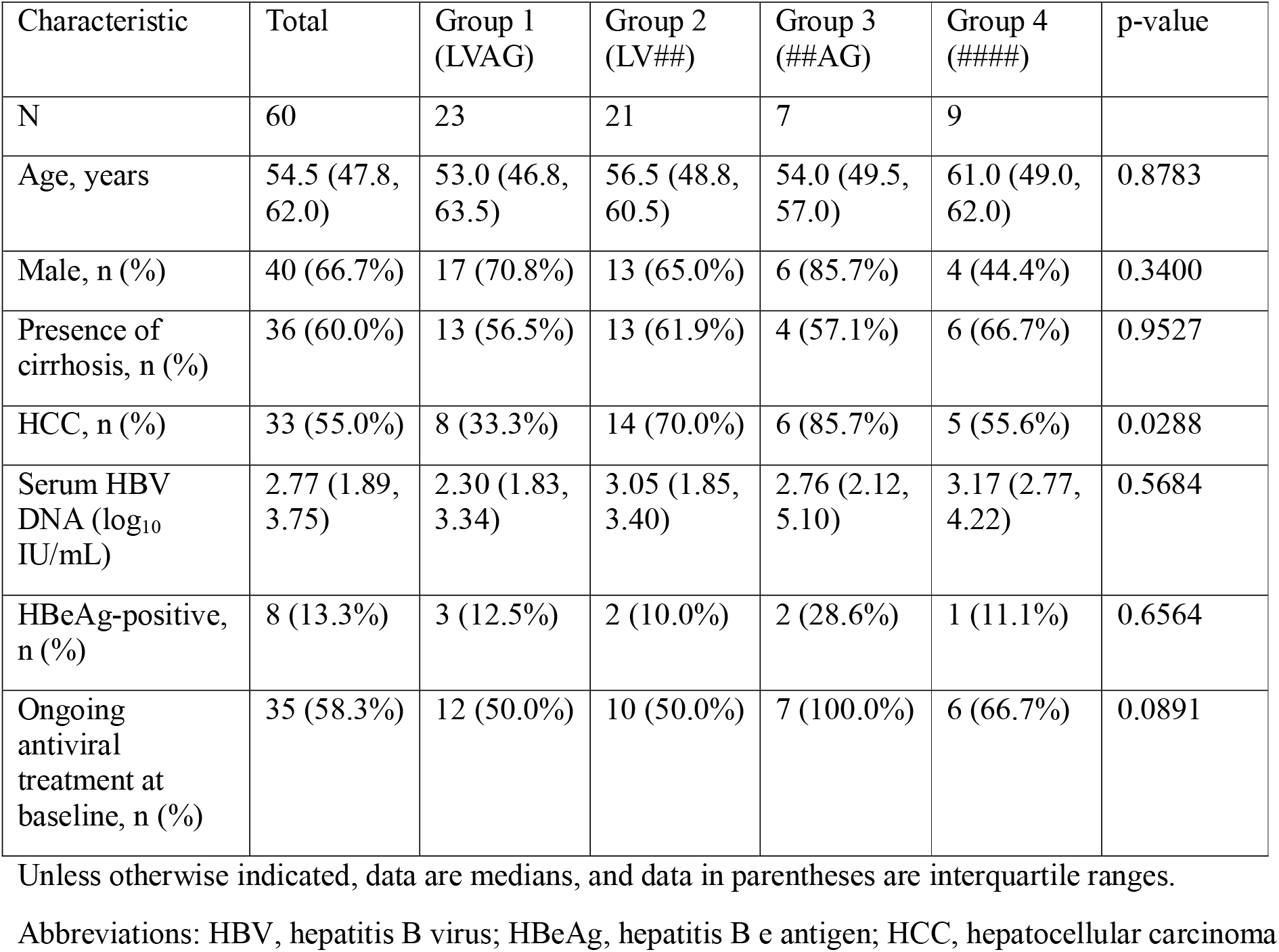
Baseline characteristics of study patients.

| Characteristic | Total | Group 1<br>(LVAG) | Group 2<br>(LV##) | Group 3<br>(##AG) | Group 4<br>(####) | p-value |
| --- | --- | --- | --- | --- | --- | --- |
| N | 60 | 23 | 21 | 7 | 9 |  |
| Age, years | 54.5 (47.8, 62.0) | 53.0 (46.8, 63.5) | 56.5 (48.8, 60.5) | 54.0 (49.5, 57.0) | 61.0 (49.0, 62.0) | 0.8783 |
| Male, n (%) | 40 (66.7%) | 17 (70.8%) | 13 (65.0%) | 6 (85.7%) | 4 (44.4%) | 0.3400 |
| Presence of cirrhosis, n (%) | 36 (60.0%) | 13 (56.5%) | 13 (61.9%) | 4 (57.1%) | 6 (66.7%) | 0.9527 |
| HCC, n (%) | 33 (55.0%) | 8 (33.3%) | 14 (70.0%) | 6 (85.7%) | 5 (55.6%) | 0.0288 |
| Serum HBV DNA (log<sub>10</sub> IU/mL) | 2.77 (1.89, 3.75) | 2.30 (1.83, 3.34) | 3.05 (1.85, 3.40) | 2.76 (2.12, 5.10) | 3.17 (2.77, 4.22) | 0.5684 |
| HBeAg-positive, n (%) | 8 (13.3%) | 3 (12.5%) | 2 (10.0%) | 2 (28.6%) | 1 (11.1%) | 0.6564 |
| Ongoing antiviral treatment at baseline, n (%) | 35 (58.3%) | 12 (50.0%) | 10 (50.0%) | 7 (100.0%) | 6 (66.7%) | 0.0891 |

Unless otherwise indicated, data are medians, and data in parentheses are interquartile ranges.
Abbreviations: HBV, hepatitis B virus; HBeAg, hepatitis B e antigen; HCC, hepatocellular carcinoma

To evaluate the changes in expression of proteins, we focused on relatively high abundant 406 protein reproducibly detected in blood. We did not restrict our analysis to HBV-related proteins. The expressional alterations were determined based on the intensities by MRM (Figure 1A). Among the four groups, the expression of 406 proteins was not significantly different (Figure 1B). (ANOVA p-value = 0.73)

**Figure. 1.** Comparative proteomic analysis of HBV genotype-associated groups. (A) Study design and grouping strategy based on predefined HBV genotype signatures. (B) Global comparison of 406 plasma proteins across the four genotype-defined groups showed no significant difference (ANOVA p = 0.73). Dot indicates individual patients.

### Comparative Proteomic Analysis of HBV Genotype-Associated Protein Expression

A total of 406 proteins were quantitatively analyzed, with statistical significance determined by adjusted p-values < 0.05. Pairwise comparisons between the four genotype groups (Group 1: LVAG, Group 2: LV##, Group 3: ##AG, Group 4: ####) demonstrated substantial heterogeneity in protein expression patterns. Group 3 exhibited the most pronounced differences in the number of significantly differentially expressed proteins compared to all other groups (Figure 1C). Specifically, the Group 3 vs Group 1 comparison yielded the highest number of differentially expressed proteins (n=56), followed by Group 3 vs Group 2 (n=39), Group 4 vs Group 1 (n=20), Group 4 vs Group 2 (n=12), Group 4 vs Group 3 (n=8), and Group 2 vs Group 1 (n=4). These findings indicate that Group 3 represents a distinct proteomic signature among the HBV genotype groups examined.

Two proteins of particular interest emerged from this analysis: CRISPLD2 (Q9H0B8) and HSPD1 (P10809). CRISPLD2 expression levels were significantly elevated in Groups 3 and 4 compared to Groups 1 and 2 (Figure 1D). Quantitative analysis revealed mean expression levels of approximately 1.0 in Groups 1 and 2, while Groups 3 and 4 showed elevated levels of 2.2 and 1.8, respectively (p < 0.001 and p < 0.05, respectively). CRISPLD2, a cysteine-rich secretory protein containing LCCL domains, functions as a critical lipopolysaccharide-binding protein involved in endotoxin regulation and inflammatory response modulation. HSPD1 (Heat Shock Protein 60) demonstrated an even more pronounced differential expression pattern, with Group 3 showing the highest expression levels, followed by Group 4, Group 2, and Group 1. The difference between Group 3 and Group 1 was highly significant (p < 0.01). Notably, HSPD1 has established direct functional interactions with HBV proteins, particularly the HBV × protein (HBx), and plays crucial roles in viral polymerase maturation and immune evasion mechanisms. Previous studies have demonstrated that HSP60 forms complexes with mitochondrial HBV proteins and contributes to viral replication efficiency through molecular chaperone functions. The concurrent upregulation of both CRISPLD2 and HSPD1 in Group 3 suggests a complex interplay between viral replication support mechanisms and host immune regulatory responses. While HSPD1 elevation may facilitate enhanced viral replication through chaperone-mediated protein folding and viral polymerase activation, increased CRISPLD2 expression indicates a compensatory anti-inflammatory response that could modulate the inflammatory cascade associated with chronic HBV infection.

These proteomic differences across HBV genotype groups provide molecular evidence for genotype-specific host-pathogen interactions and suggest that therapeutic approaches may need to be tailored according to the underlying genotype-associated protein expression profiles. The identification of these differentially expressed proteins offers potential biomarkers for genotype-specific disease progression and treatment response prediction in chronic hepatitis B patients.

### Co-expressed proteins depending mutation group

To investigate coordinated protein expression patterns across the four groups, we applied weighted gene co-expression network analysis (WGCNA) to the 406 proteins quantified by LC–MS. A total of eight distinct protein modules were identified (Figure 2A). Correlation analysis between module eigengenes and group comparisons revealed that three modules (Modules 5, 6, and 7) were strongly associated with group differences. Module 5 consisted of 31 proteins, Module 6 of 57 proteins, and Module 7 of 111 proteins. These three modules showed significant correlations with contrasts involving Group 3 (##AG). For example, Module 6 was positively correlated with Group 2 vs. Group 3 (r = 0.67, p < 0.001) and Group 3 vs. Group 4 (r = 0.59, p < 0.05), while Module 7 showed a similar pattern (Group 2 vs. Group 3: r = 0.57, p < 0.001; Group 3 vs. Group 4: r = 0.47, p < 0.01). Module 5, in contrast, was negatively correlated with Group 2 vs. Group 3 (r = –0.51, p < 0.01) but positively correlated with Group 1 vs. Group 4 (r = 0.45, p < 0.01). These results indicate that multiple co-regulated protein modules distinguish Group 3 from the other groups. Analysis of module eigengenes further confirmed these associations. Eigenprotein values of Modules 5, 6, and 7 were significantly different across the four groups (ANOVA p = 0.0036, 0.0015, and 0.0056, respectively) (Figure 2C). In all three modules, Group 3 consistently exhibited higher eigenprotein values compared with the other groups, with post hoc comparisons showing significant differences particularly between Group 3 and Groups 1 or 2. This consistent elevation suggests that proteins in these modules are coordinately upregulated in Group 3.

**Figure.2.** Differentially expressed proteins and co-expression analysis. (A) Pairwise comparisons between the four groups revealed heterogeneous proteomic alterations, with Group 3 (##AG) showing the largest number of differentially expressed proteins. (B) Differential expression of CRISPLD2 and HSPD1 across groups, demonstrating significant upregulation particularly in Group 3.

To further dissect the heterogeneity within each module, we performed k-means clustering of individual protein expression patterns. This analysis revealed distinct subclusters of proteins within Modules 5, 6, and 7 that shared highly similar expression trajectories across the four groups (Figure 2D). Clustering was performed based on expression levels despite similar overall patterns. Group 3 showed the highest expression across modules, except for Module 5 where Group 4 had the greatest increase.

By combining module-level and cluster-level analyses, we identified not only broad co-regulated protein signatures but also the subsets of proteins that may drive the observed group-specific changes. These results highlight that the distinct protein expression patterns in Group 3 are not uniformly distributed across all proteins in a module, but instead are concentrated within specific clusters of proteins, pointing to potentially critical subsets that underlie the transition between [LV] and [##].

### Network Analysis of Protein Modules and PI3K/Akt Pathway in HBV Infection

Network analysis of proteins within modules 5, 6, and 7 revealed distinct functional clustering patterns with significant biological relevance to HBV pathogenesis (Figure 3A). The network topology demonstrated that these modules exhibited high interconnectivity score (clustering coefficient > 0.7), indicating tightly regulated protein complexes rather than randomly associated proteins. Modules 5, 6, and 7 demonstrated pronounced enrichment in focal adhesion pathways was significantly enriched (p<0.001). Beyond focal adhesion, these modules showed significant enrichment in extracellular matrix organization (p<0.001), angiogenesis regulation (p<0.001), and cell migration pathway (p<0.001), collectively pointing toward a coordinated response involving tissue remodeling and vascular adaptation during chronic HBV infection. The co-occurrence of these pathways within the same modules suggests functional synergy between cellular adhesion, matrix degradation, and neovascularization processes.

**Figure. 3.** Functional enrichment of co-expressed protein modules. (A) Weighted gene co-expression network analysis (WGCNA) identified eight protein modules, with Modules 5, 6, and 7 showing the strongest associations with group differences. (B) Module eigengene correlation with genotype group contrasts revealed significant associations with focal adhesion, extracellular matrix organization, angiogenesis, and cell migration. (C) Line plots showing subgrouped protein expression trajectories for (left) Module 5, (middle) Module 6, and (right) Module 7 across the four genotype-defined groups. Proteins within each module were partitioned into optimal clusters by k-means clustering, with individual protein expression profiles shown as thin lines and cluster mean trajectories highlighted as bold lines.

Most notably, PI3K-Akt signaling pathway components (hsa04151 and WP4172) were prominently enriched within these protein modules (p<0.001) (Figure 3B). The PI3K/Akt pathway plays a critical dual role in HBV infection, where HBx directly activates PI3K/Akt signaling in hepatocytes (24). This activation paradoxically suppresses HBV replication by downregulating hepatocyte nuclear factor 4α (HNF4α) while simultaneously promoting hepatocyte survival through apoptosis inhibition. Treatment with PI3K inhibitors significantly enhances HBV replication, confirming the pathway’s role as a negative regulator of viral replication (25). The pathway also contributes to HBV-associated oncogenesis through HBx phosphorylation by Akt1 kinase and alpha-fetoprotein upregulation, creating a positive feedback loop that promotes hepatocarcinogenesis (26). The network connectivity among modules 5, 6, and 7, particularly the co-enrichment of angiogenesis functions with PI3K/Akt signaling, suggests coordinated regulation of pathways involved in tumor microenvironment modification during chronic HBV infection.

### Protein-Protein Interaction Enrichment Analysis of High-Density Network Clusters

We focused on high-density interaction regions within the protein network to identify functionally coherent subnetworks, the used proteins were obtained from module 5, 6, and 7. Network topology analysis revealed four distinct protein clusters with varying connectivity patterns and functional specialization (Figure 4, upper panel). The largest cluster (red, n=23) demonstrated the highest intracluster connectivity. Hub protein identification revealed key regulatory nodes that serve as critical bridges between functional modules.

**Figure. 4.** Protein–protein interaction (PPI) enrichment analysis of high-density clusters. High-density interaction subnetworks derived from Modules 5, 6, and 7 revealed four functional clusters. The largest cluster (red, n=23) showed the highest intracluster connectivity. Functional enrichment highlighted pathways related to protein folding, chaperone-mediated complex assembly, extracellular matrix interactions, and cytoskeletal organization.

**Figure. 5.** Network topology and functional specialization of densely connected protein clusters. Protein–protein interaction network analysis identified four clusters with distinct functional enrichment patterns, including chaperone-mediated protein complex assembly, protein folding, extracellular matrix organization, and cytoskeletal regulation. **N**ode size represents the average magnitude of expression differences across genotype groups, and statistically significant differences are highlighted in yellow.

Protein-protein interaction enrichment analysis of these densely connected clusters revealed several critical biological processes with high statistical significance (Figure 4). The most prominent enriched functions included chaperone-mediated protein complex assembly (GO:0051131, -log p-value = 9.0), protein folding (GO:0006457, -log p-value = 6.6), indicating severe disruption of cellular protein homeostasis during HBV infection. The chaperone network cluster, primarily composed of heat shock proteins (HSPA4, HSPD1) and protein disulfide isomerase family members (PDIA3, PDIA6), suggests activation of the unfolded protein response (UPR) pathway (19). This cellular stress response is commonly triggered by HBV surface protein accumulation in the ER, leading to ER stress and subsequent chaperone upregulation as a protective mechanism. Cytoskeleton organization in muscle cells (hsa04820, -log p-value = 10.6), ECM proteoglycans (R-HSA-3000178, -log p-value = 10.0), and integrin cell surface interactions (R-HSA-216083, -log p-value = 9.9). These findings indicate that HBV infection significantly impacts protein quality control mechanisms, cellular structural maintenance, and extracellular matrix interactions, suggesting widespread perturbation of fundamental cellular processes during viral pathogenesis (19, 27, 28). The coordinate dysregulation of these pathways creates a cellular environment conductive to chronic inflammation, progressive fibrosis, and eventually malignant transformation. Notably, the pathway enrichment pattern observed here closely resembles that seen in HCC, suggesting that these proteomic changes may serve as early molecular markers for cancer risk stratification in chronic hepatitis B patients. The high statistical significance of these functional categories (all p-values < 0.001) and their interconnected nature suggest that therapeutic interventions targeting multiple pathways simultaneously may be more effective than single-pathway approaches in managing HBV-induced liver disease progression.

## Conclusions

In conclusion, genotype-informed stratification of 60 HBV-infected patients revealed a coherent, systems-level host response that was not apparent in the global comparison of all four groups (ANOVA across 406 plasma proteins, p=0.73), but became evident in pairwise contrasts and network-level analyses. The ##AG group—harboring the basal core promoter/precore signature in the absence of the mutation related to liver disease progression LV motif—exhibited the most distinctive proteomic phenotype, including marked elevations of CRISPLD2 and HSPD1, implicating a coupled axis of anti-inflammatory buffering and chaperone-assisted viral processes. Convergent approaches (WGCNA, k-means subclustering, and protein–protein interaction enrichment) identified tightly connected modules (high clustering coefficients) that differentiate this group and are enriched for focal adhesion, extracellular matrix organization, angiogenesis, and cell migration, alongside prominent PI3K/Akt signaling. This pathway aligns with known HBV biology—where PI3K/Akt activation restrains replication while promoting hepatocyte survival—and maps onto microenvironmental remodeling processes linked to progression toward hepatocarcinogenesis. Together, these data provide molecular evidence for genotype-specific host–pathogen interactions in chronic HBV infection, nominate CRISPLD2 and HSPD1 as accessible biomarker candidates, and argue for therapeutic strategies that co-target PI3K/Akt and microenvironmental pathways rather than single node. Although derived from a moderate-sized cohort and a panel restricted to high-abundance plasma proteins, the findings demonstrate internal consistency across orthogonal analyses, supporting their robustness and warranting validation in independent and longitudinal cohorts to refine genotype-tailored risk stratification and treatment.

## Acknowledgement

This work was supported by the National Research Foundation of Korea (NRF) grant funded by the Korean government (MSIT) (RS-2023-00209456) and the Korea Basic Science Institute (National Research Facilities and Equipment Center) grant funded by the Korean government (MSIT) (RS-2024-00402298). This research was also supported by the National Research Foundation of Korea (NRF), funded by the Ministry of Education (RS-2025–00553721). We acknowledge the use of Claude Opus 4.1 solely for linguistic refinement and grammatical corrections in manuscript preparation. All scientific content, data analysis, and intellectual contributions presented herein were developed independently by the authors without the use of generative AI tools. Figure 1 was created with BioRender.com (accessed 1 July 2025).

The biospecimens for this study were provided by the Seoul National University Hospital Human Biobank, a member of the Korea Biobank Network, which is supported by the Ministry of Health and Welfare.

