## Supplementary material for "Proteomic Signatures of Hepatitis B Virus Mutations Reveal Genotype-Specific Host Responses and Biomarker Candidates": Figures

### Slide 1
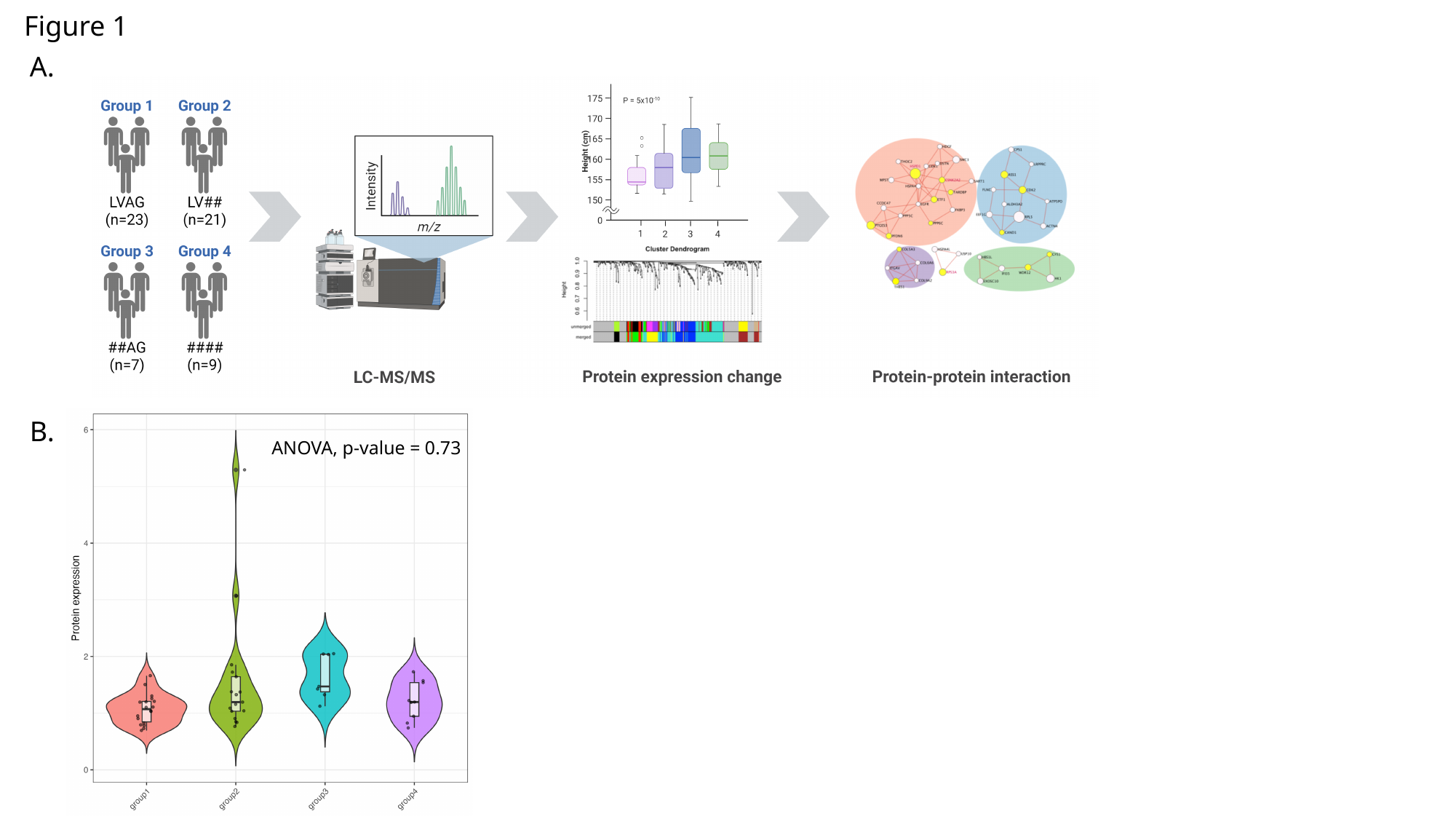

Figure 1
A.
ANOVA, p-value = 0.73
B.

### Slide 2
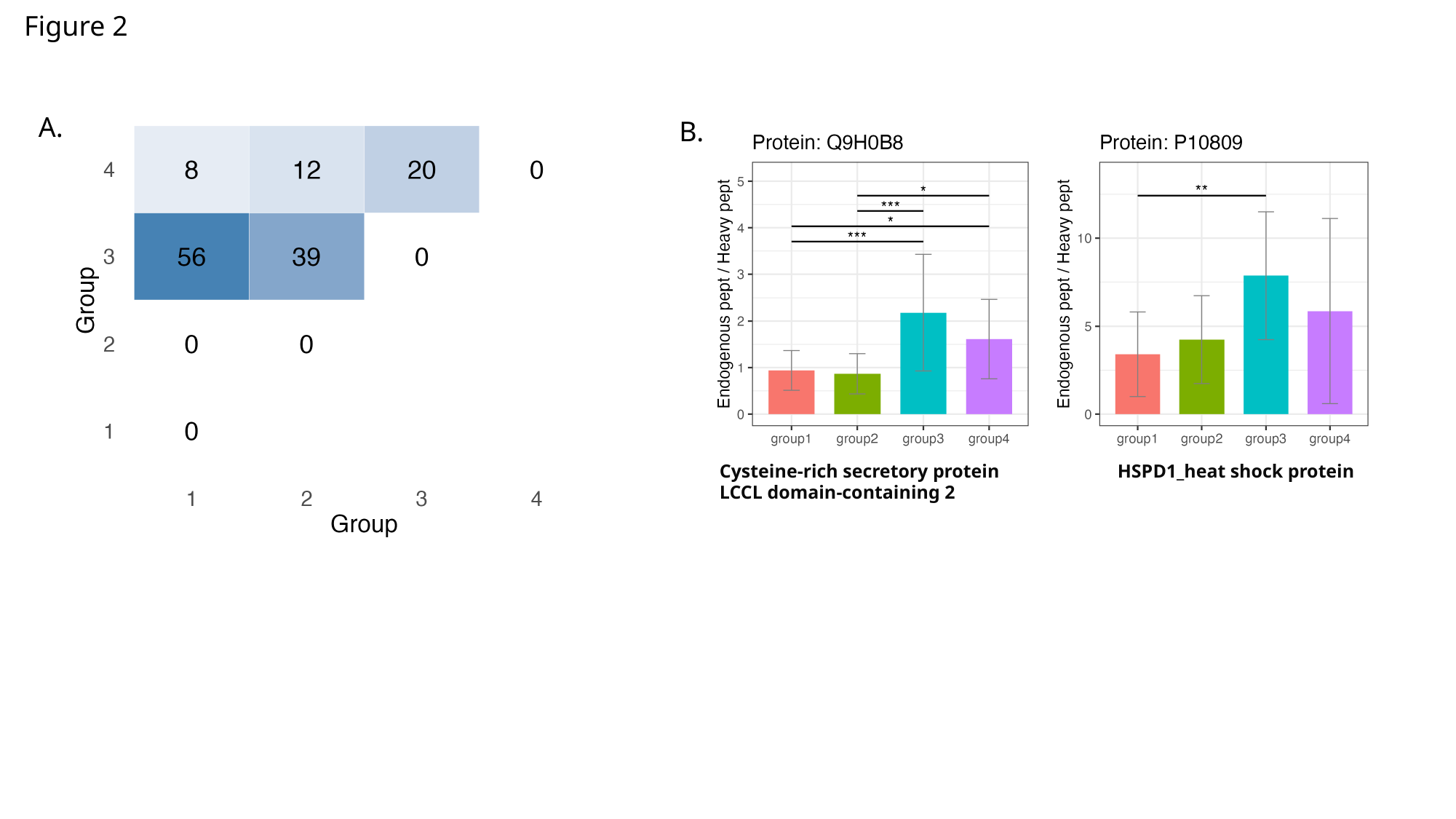

Figure 2
A.
B.
Cysteine-rich secretory protein LCCL domain-containing 2
HSPD1_heat shock protein

### Slide 3
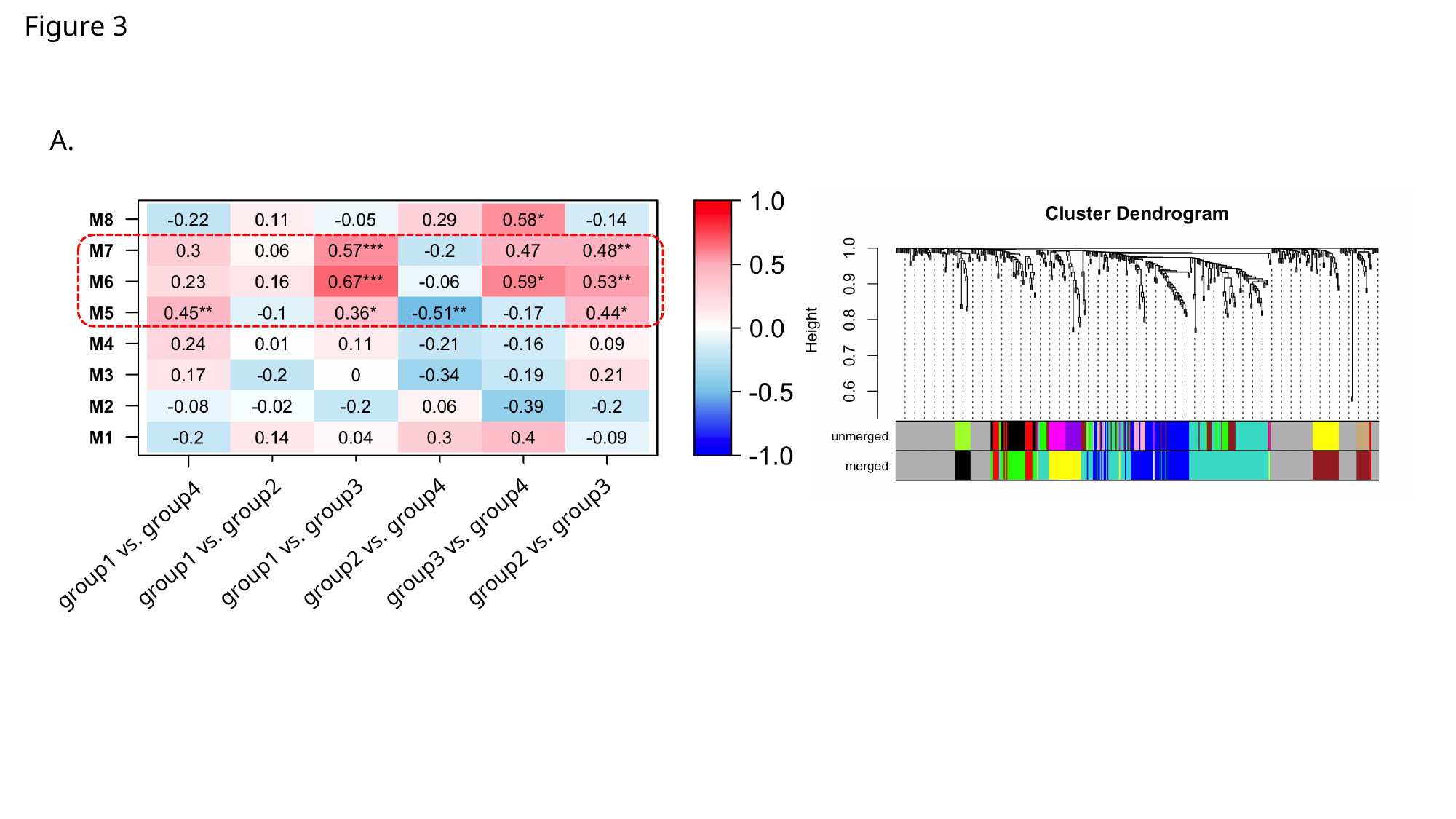

Figure 3
A.
group1 vs. group4
group1 vs. group2
group1 vs. group3
group2 vs. group4
group3 vs. group4
group2 vs. group3

### Slide 4
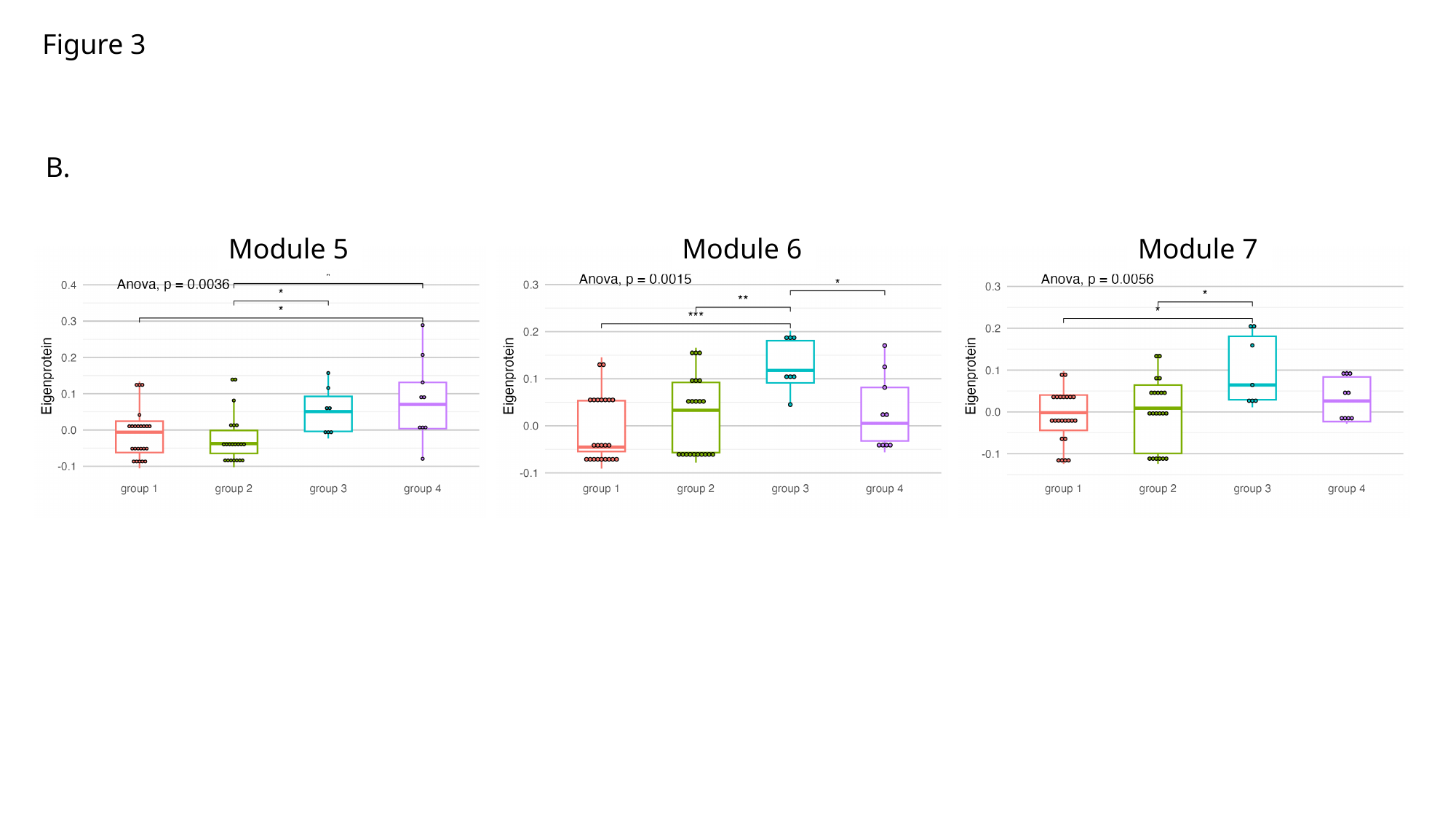

Figure 3
B.
Module 5
Module 6
Module 7

### Slide 5
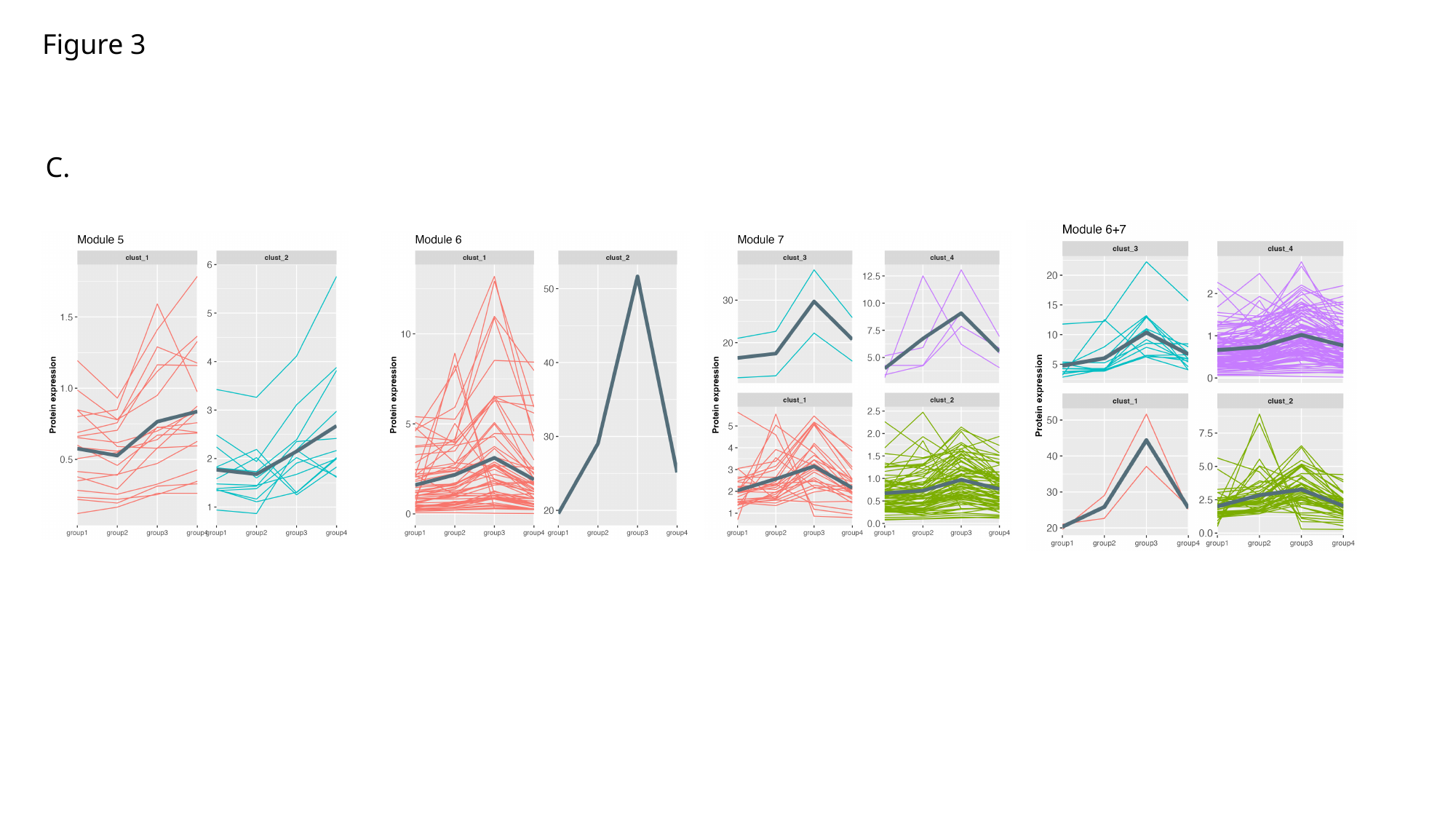

Figure 3
C.

### Slide 6
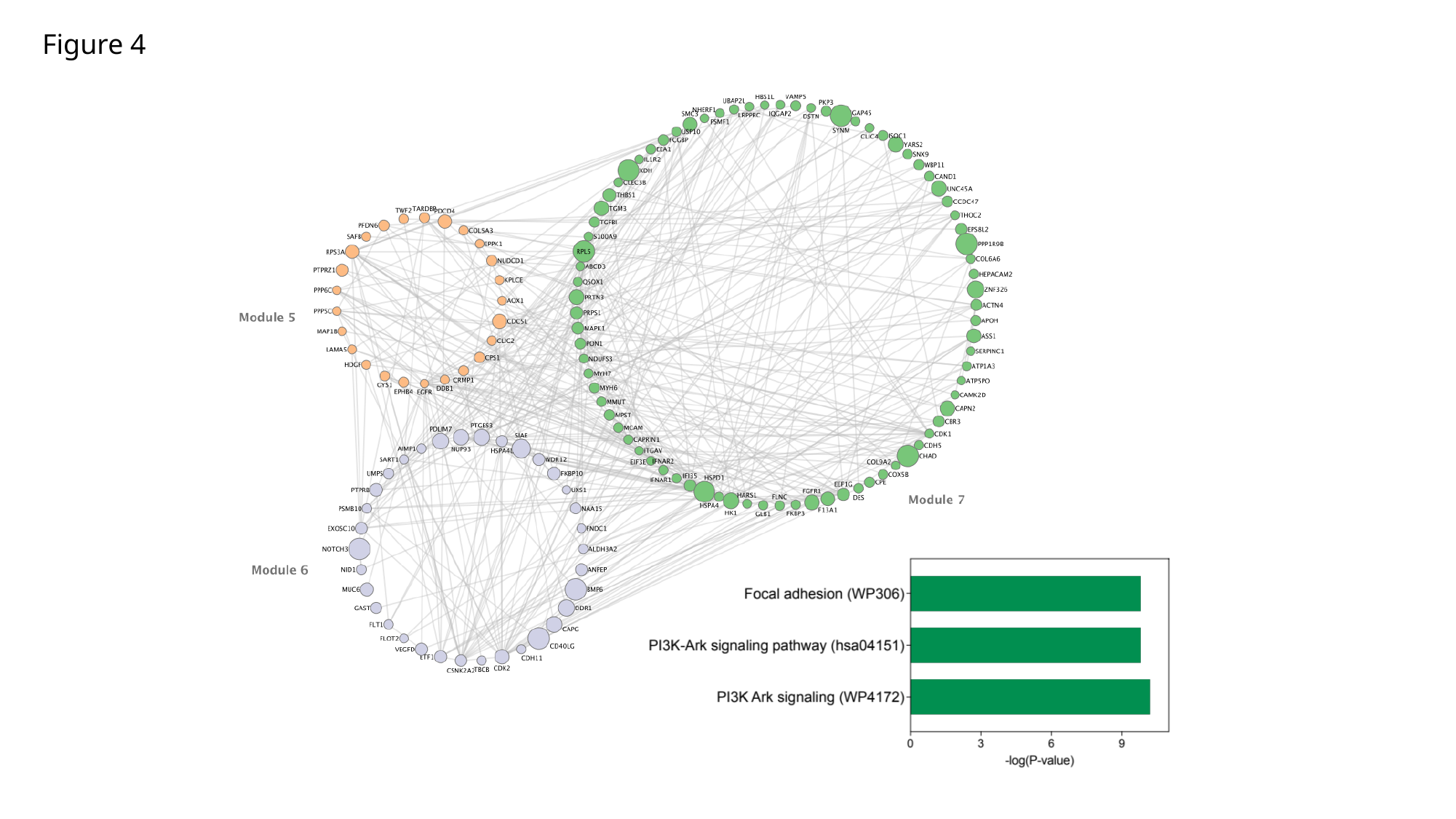

Figure 4

### Slide 7
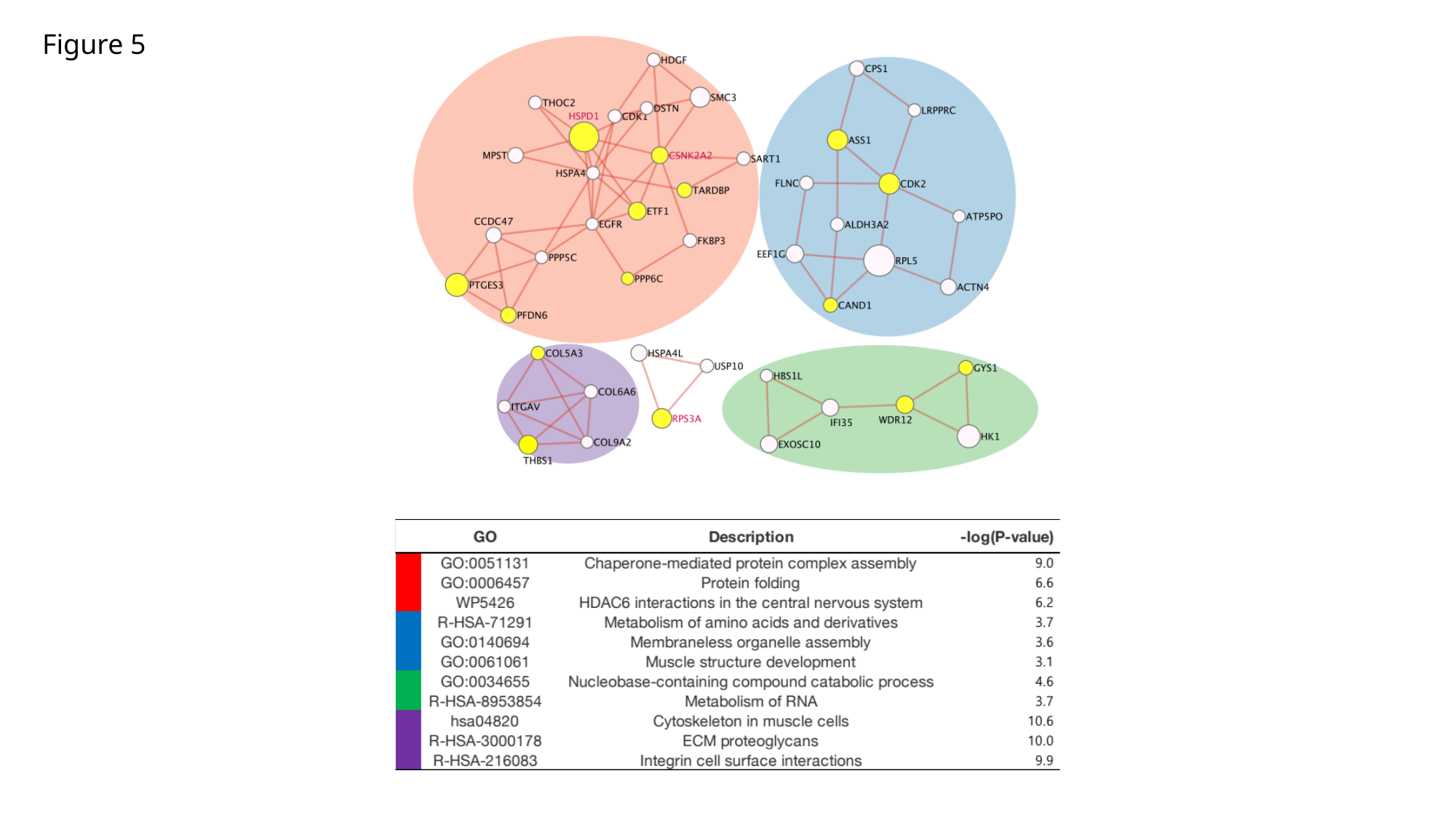

Figure 5
